# A palmitoyl transferase chemical genetic system to map ZDHHC-specific *S*-acylation

**DOI:** 10.1101/2023.04.18.537386

**Authors:** Cory A. Ocasio, Marc P. Baggelaar, James Sipthorp, Ana Losada de la Lastra, Jana Volarić, Christelle Soudy, Elisabeth M. Storck, Susana A. Palma-Duran, James I. MacRae, Goran Tomic, Lotte Carr, Julian Downward, Ulrike S. Eggert, Edward W. Tate

## Abstract

The 23 human ZDHHC *S*-acyltransferases catalyze long-chain *S*-acylation at cysteine residues across an extensive network of hundreds of proteins important for normal physiology or dysregulated in disease. Here we present a technology platform to directly map the protein substrates of a specific ZDHHC for the first time at the whole proteome level, in intact cells. Structure-guided engineering of paired ZDHHC ‘hole’ mutants and ‘bumped’ chemically tagged fatty acid probes enabled probe transfer to specific protein substrates with excellent selectivity over wild type ZDHHCs. Chemical genetic systems were exemplified for five ZDHHCs (3, 7, 11, 15 and 20), and applied to generate the first *de novo* ZDHHC substrate profiles, identifying >300 unique and shared substrates across multiple cell lines and *S*-acylation sites for novel functionally diverse substrates. We expect that this powerful and versatile platform will open a new window on *S*-acylation biology for a wide range of models and organisms.

## Introduction

The chemical and functional diversity of proteins encoded by the human genome is expanded by orders of magnitude through post-translational modification (PTM)^1, 2^. Long chain *S*-acylation is among the most widespread PTMs mediated in all eukaryotes by the ZDHHC *S*-acyltransferase family of integral membrane enzymes, including 23 known human ZDHHCs which acylate >3000 cysteine residues across ca. 12% of the human proteome^3–6^. The two-stage ZDHHC catalytic cycle commences with auto-*S*-acylation of a conserved Asp-His-His-Cys (DHHC) by C14:0 to C22:0 acyl-CoA (commonly palmitoyl (C16:0)-CoA) with concomitant release of CoASH, followed by *S*-acyl transfer to a substrate protein cysteine positioned proximal to the ZDHHC catalytic site (Fig. 1A-B)^7–9^. Protein substrates do not exhibit a clear consensus sequence beyond the requirement for a free cysteine^10^, and substrate recruitment mechanisms may include co-localization through protein interactions, membrane associated domains, or prior lipid PTMs^5, 11–13^. *S*-acylation increases local hydrophobicity and membrane affinity, and can regulate membrane microdomain partitioning, protein stability, trafficking, nuclear localization, secretion, or protein-protein interactions^14, 15^. De-*S*-acylation by acyl-protein thioesterases (Fig. 1A) creates a dynamic *S*-acylation cycle implicated in signaling cascades ^16–18^, with numerous examples of up- or down-regulation of *S-*acylation promoting pathological conditions including cancer, inflammatory disease or neurodegeneration^4, 19–22^.

**Figure 1.**
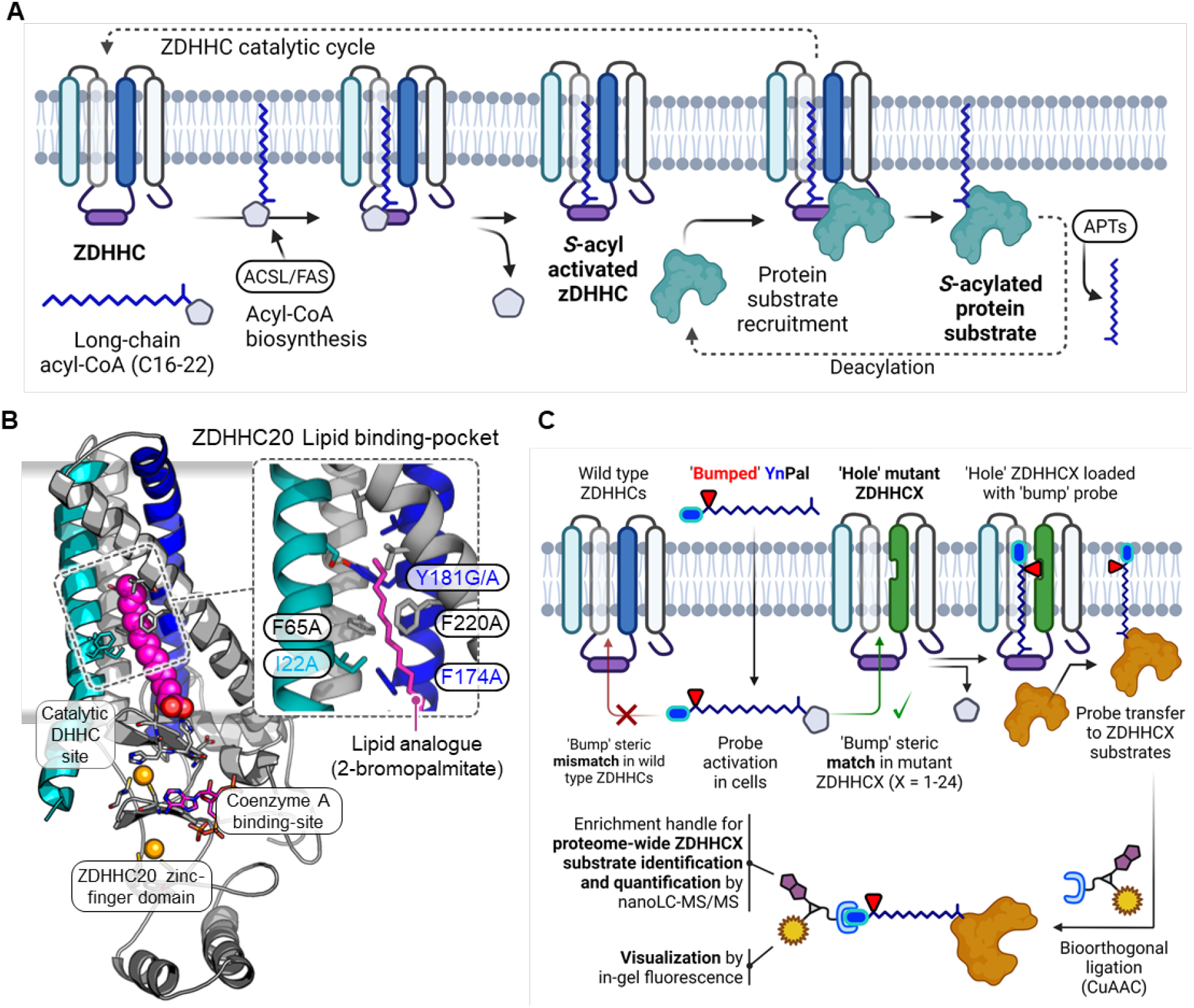
ZDHHC chemical genetics. (**A**) *S*-acylation is mediated by ZDHHC loading of long-chain acyl-CoA derived from lipid biosynthesis followed by acyl transfer to a proximal Cys of a protein substrate and regeneration of apo-ZDHHC; the reversible cycle is closed by acyl protein thioesterase (APTs) thioester hydrolysis. (**B**) X-ray structure of human ZDHHC20 irreversibly inhibited by lipid mimic 2-bromopalmitate (PDB entry 6BML^3^); inset: sterically demanding residues in the ZDHHC20 lipid binding pocket contact the acyl chain distal to the DHHC catalytic site. (**C**) Steric complementation between a ZDHHC ‘hole’ mutant and an alkyne-tagged ‘bumped’ lipid substrate probe enables selective loading and tag transfer to ZDHHC substrates, bypassing endogenous (wild type) ZDHHCs; fluorescence visualization and chemical proteomics is enabled by bioorthogonal conjugation to multifunctional capture reagents.

Despite the importance of ZDHHCs to human health and disease, mapping the substrate network of a given ZDHHC remains a formidable challenge. Global enrichment of *S*-acylated proteins through metabolic labeling with alkyne-tagged lipid analogues or chemical exchange of *S*-acyl thioesters for affinity tags can circumvent the notorious problems of lability, poor ionization and low solubility of native *S*-acylated peptides in liquid chromatography-mass spectrometry (LC-MS), and has revolutionized our view of *S*-acylation by generation of large databases of putative substrate proteins^6, 23, 24^. However, there is currently no approach to directly associate a specific ZDHHC with cognate *S*-acylated protein substrates, and this endeavor is further complicated by the current lack of potent and selective ZDHHC inhibitors^25, 26^, and the confounding influence of ZDHHC overexpression, knockdown or knockout which can lead to redundancy, compensation and loss of ZDHHC protein interactions or ZDHHC co-regulation^27–30^.

Here, we establish the first chemical genetic system for direct labeling and identification of the substrates of a specific ZDHHC in intact cells, through steric complementation (so-called ‘bump and hole’) (Fig. 1C)^31–33^. We report mutant/probe pairs for five diverse ZDHHCs (ZDHHCs 3, 7, 11, 15 and 20) and demonstrate mutant-specific ZDHHC-loading and protein substrate transfer with high selectivity over wild type ZDHHCs. Coupled to chemical proteomics, this technology enabled *de novo* identification of >300 ZDHHC-specific substrates and *S*-acylation sites in varied human cell lines, and the first extended substrate networks for ZDHHCs 7, 15 and 20. Adaptability and ease of implementation across cellular models suggests that ZDHHC chemical genetics offers a novel platform for systematic investigation of ZDHHC biology, with the potential to catalyze knowledge-driven selection of ZDHHCs and ZDHHC-mediated pathways for future therapeutic validation or biomarker discovery.

## Results

### Selective *S*-acylation by an engineered ZDHHC20 mutant/probe pair

Steric complementation for ZDHHCs imposes stringent requirements on mutant and probe design: the mutant ZDHHC should retain wild type activity and protein substrate specificity; the probe must bear both a ‘bump’ and an alkyne tag, and be efficiently activated to the CoA thioester form in the cell without interfering with endogenous lipid metabolism; and the activated probe must be minimally processed by wild type ZDHHCs to deliver a selectivity window for ZDHHC-specific substrate identification. We therefore focused on mutations and probe ‘bump’ modifications distal to the DHHC active site to minimize interference with catalytic activity or lipid probe activation. Human ZDHHC20 crystal structures reveal a conical transmembrane lipid-binding pocket in a four-pass transmembrane (4TM) helix domain adjacent to the cytosolic catalytic site^3, 34^, and this structure was used as a template to design six ZDHHC20 mutations towards the apex of the 4TM domain with potential for steric complementation with a matching ‘bumped’ lipid probe (Fig. 1C). A panel of nine alkyne-tagged bumped lipid analogs were designed and synthesized, positioning small (acetyl, Ac), medium (cyclopropanecarbonyl, *c*Pr) or large (benzoyl, Bz) bump groups at increasing distance from the acid (Fig. 2A, Methods). These probes together encompass the most common linear chain lengths found in endogenous *S*-acylation (16, 18 or 20 atoms) ^8^, enabling pairing to mutants with differing ‘hole’ size and position, whilst he alkyne tag permits ligation of fluorescent reporters and/or affinity handles to modified proteins through copper-catalyzed alkyne-azide cycloaddition (CuAAC) (Fig. 1C), revealing ZDHHC autoacylation and substrates in cellular assays. Probe optimization was envisaged as a two-stage process, first determining ideal chain length and bump placement, followed by screening the selected chain-length for optimal bump size (Fig. 2B).

**Figure 2.**
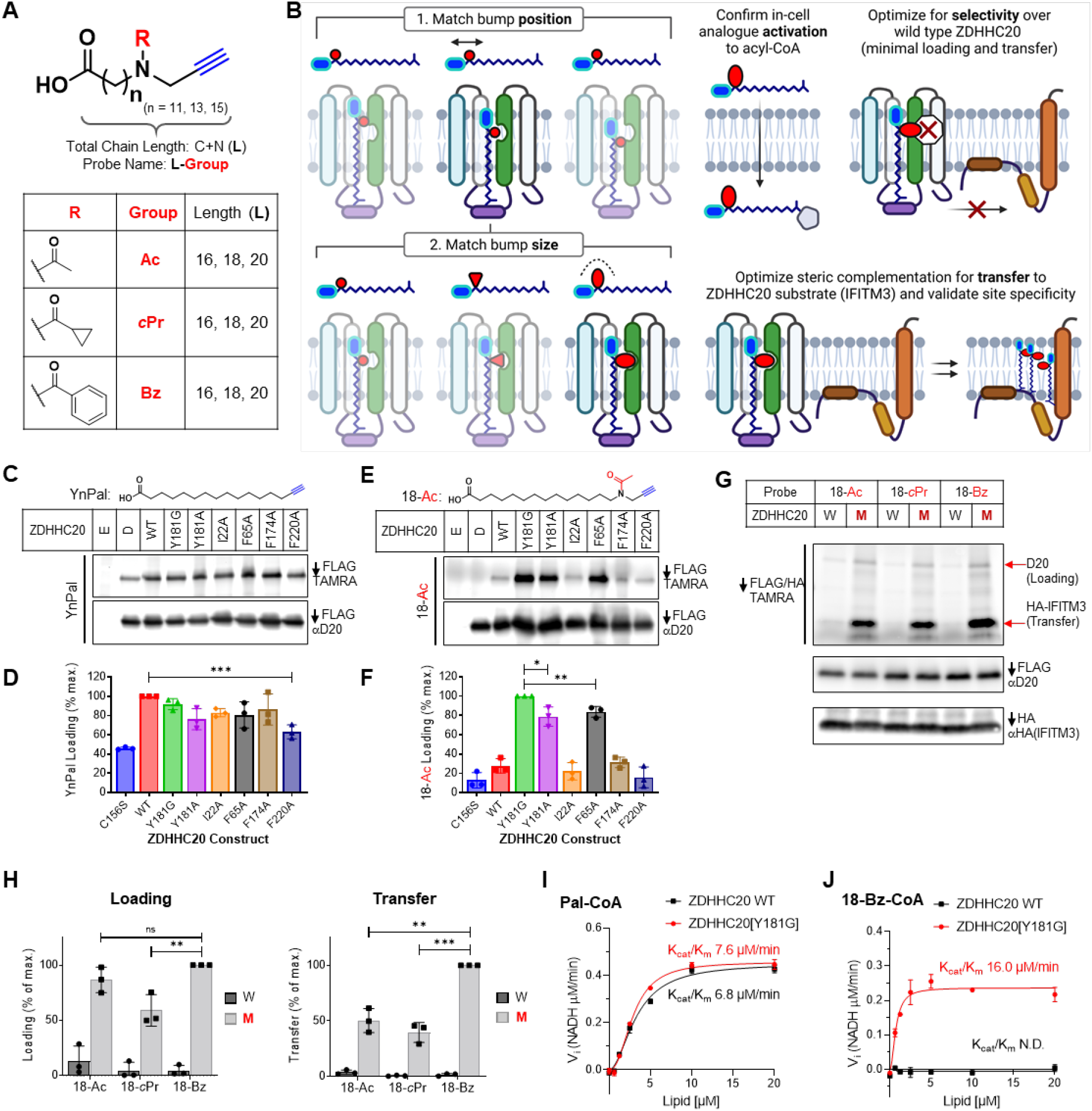
Engineering a ‘bump’ probe and ‘hole’ mutant pair for ZDHHC20. (**A**) Fatty acid probes containing an alkynyl click-handle (blue), varying chain-length **L** = 16, 18 or 20 heavy atoms in the chain (carbons + nitrogen), and a R ‘bump’ group (red): acetyl (Ac), cyclopropanecarbonyl (*c*Pr) or benzoyl (Bz). (**B**) Two-stage pairing strategy for a designed ZDHHC20 mutant optimizes probe chain length then bump size to match the new binding cavity, with probe activation, selectivity over ZDHHC20 wild type and transfer to a known ZDHHC20 substrate (IFITM3) optimized in parallel. (**C-F**) Bump-hole loading analysis of C-terminal FLAG-tagged ZDHHC20 WT and mutants in HEK293T cells treated with 15 μM YnPal (**C** & **D**) or 18-Ac (**E** & **F**) for 4 h (E = empty vector, D = catalytic-dead ZDHHC20[C156S]) (n=3). (**G**) Probe bump-size optimization by transfer assays with HA-IFITM3 and either WT ZDHHC20 (W) or ZDHHC20[Y181G] (M) co-expression in HEK293T cells (n=3). (**H**) Average loading and transfer activity relative to highest fluorescent/input ratio (n = 3 ± S.D). (**I-J**) Enzyme kinetics for WT ZDHHC20 and ZDHHC20[Y181G] treated with (**I**) Pal-CoA or (**J**) 18-Bz-CoA using a KDH assay (*3*). Michaelis-Menten plots generated from average reaction rate (n = 3) (NADH generated (μM)/min) ± S.D. vs. lipid concentration (μM) using Prism 9.0. P-values determined using Prism 9.0 unpaired t-test statistics module (ns (not statistically significant), P > 0.05; *, P ≤ 0.05; **, P ≤ 0.01; ***, P ≤ 0.001 and ****, P ≤ 0.0001.

We first explored baseline autoacylation activity for each ZDHHC20 mutant by metabolic labeling with YnPal (an alkyne-tagged analog of palmitate, C16:0) in HEK293T cells expressing each of six FLAG-tagged ZDHHC20 mutants (Fig. 2C-D)^23^. FLAG immunoprecipitation followed by CuAAC ligation to TAMRA and analysis by in-gel fluorescence confirmed autoacylation activity for all mutants, with ZDHHC20[Y181G] showing labeling equivalent to wild type (WT). Residual acylation was observed for catalytically dead Cys mutant ZDHHC20[C156S] (lane ‘D’, Fig. 2C-D, Extended Data Fig. 1A-B), consistent with previously characterized *S*-acylation at non-catalytic ZDHHC20 Cys residues mediated by endogenous ZDHHCs^24, 35^. As expected, labeling was sensitive to thioester cleavage by hydroxylamine (HA), and increased with YnPal concentration and incubation time, leading to steady state labeling after 2 hours incubation with 15 µM YnPal (Extended Data Fig. 1A-H). An initial screen of ZDHHC20 mutants under similar conditions against bumped probe 18-Ac (18-atom chain length, smallest bump) revealed strong complementation for F65 and Y181 mutants and reduced loading with WT ZDHHC20, with Y181G exhibiting 5-fold higher loading than WT (Fig. 2E-F). Furthermore, residual ZDHHC20[C156S] labeling was suppressed to background, suggesting that the bumped probe is a poor substrate for endogenous ZDHHCs which *S*-acylate ZDHHC20 *in trans*. These data are consistent with previous evidence that mutations in the 4TM domain can tolerate longer chain lipids^3, 8^, and encouraged us to proceed to optimize steric complementation with ZDHHC20[Y181G].

We implemented a structure-guided screening sequence commencing with the smallest ‘bump’ probes of increasing chain length (16-Ac, 18-Ac or 20-Ac) to identify the length register matching the bump to the mutant cavity (Fig. 2B, Extended Data Fig. 2A-B). An integrated assay format was used to assess both lipid loading and transfer to substrate by co-expressing ZDHHC20-FLAG (WT or Y181G) with a canonical ZDHHC20 substrate, HA-tagged IFITM3^21, 36^, enabling sensitive in-gel fluorescence quantification of ZDHHC20 and protein substrate labeling following dual FLAG/HA pull-down and on-bead CuAAC ligation to TAMRA-azide. 18-Ac and 20-Ac were similarly superior to 16-Ac in ZDHHC20[Y181G] loading, but 18-Ac transfer to IFITM3 was twofold higher than 20-Ac (Extended Data Fig. 2C-D), implying improved catalytic efficiency. Bump size screening (Ac, *c*Pr or Bz) at the 18-atom chain length identified 18-Bz as an optimal probe for ZDHHC20[Y181G], exhibiting >20-fold higher loading and >60-fold more efficient transfer than WT ZDHHC20 (Fig. 2G-H).

### Steric complementation delivers catalytic efficiency orthogonal to wild type ZDHHC20

We next compared the enzyme kinetic parameters of activated 18-Bz CoA thioester (18-Bz-CoA) and YnPal-CoA for recombinant FLAG-purified WT ZDHHC20 or ZDHHC20[Y181G] using an established real-time enzyme-coupled assay, measuring CoA generation during spontaneous turnover of auto-*S*-acylated ZDHHC20 in the absence of a protein substrate (Extended Data Fig. 3)^3, 37^. Consistent with cellular assay data, YnPal-CoA had similar catalytic efficiency (k_cat_/K_M_) for WT and ZDHHC20[Y181G] (6.8 ± 0.3 and 7.6 ± 0.3 μM/min, respectively) (Fig. 2I, Extended Data Table 1). Furthermore, 18-Bz-CoA had even greater catalytic efficiency with ZDHHC20[Y181G] (16.0 ± 1.0 μM/min), with slightly reduced k_cat_ and K_M_ relative to YnPal-CoA, whilst showing no measurable activity with WT ZDHHC20 (Fig. 2J and Extended Data Table 1). As expected, catalytically dead ZDHHC20[C156S] and [Y181G/C156S] mutants were inactive in this assay (Extended Data Fig. 3H-I). Together, these data provide compelling biochemical evidence that designed ZDHHC steric complementation can deliver orthogonal ZDHHC loading at a level comparable to wild type.

**Figure 3.**
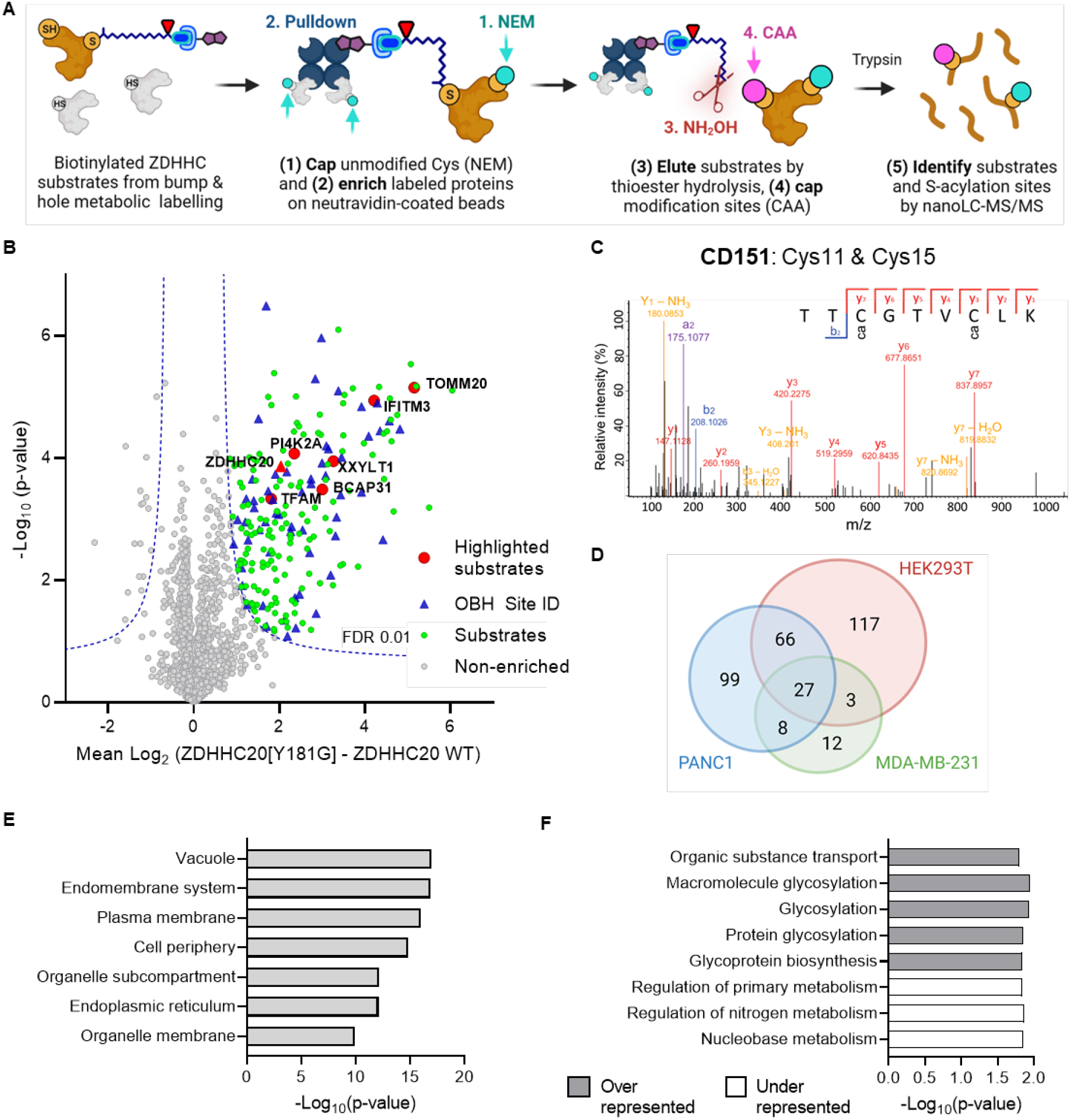
Chemical proteomic ZDHHC20 substrate and modification site identification. **(A)** Chemical proteomic on-bead hydrolysis (OBH) workflow for enrichment and identification of *S*-acyltransferase substrates and *S*-acylation sites by LC-MS/MS. (**B**) Chemical proteomic analysis of ZDHHC20 substrates in HEK293T cells (15 µM 18-Bz, 8 h); enrichment in ZDHHC20[Y181G] cells over WT ZDHHC20 reveals selective ZDHHC20 loading (red triangle), and significantly enriched substrates (green circles) selected for further validation (red circles), and with site identification data (blue triangles) (Student’s unpaired T-test, S_0_ 0.5, FDR 0.01, n=4 per condition). (**C**) LC-MS/MS spectrum corroborating reported sites of CD151 *S*-acylation at Cys11 and Cys15 (see also Extended Data Figure 9). (**D**) Venn diagram of chemical genetic ZDHHC20 substrate maps in HEK293T, MDA-MB231 and PANC1 cells. (**E**, **F**) Statistical overrepresentation analysis for chemical genetic ZDHHC20 substrates for (**E**) cellular compartment (Slim)-GO terms compared to the human proteome showing terms with -Log_10_(p-value) >9 from an FDR adjusted Fisher’s exact test and (**F**) biological process GO-terms compared to a reference database of all known *S*-acylated proteins (SwissPalm), using PANTHER classification showing terms with -Log_10_(p-value) >1.5 from an FDR adjusted Fisher’s exact test.

### Chemical genetics enables ZDHHC20-specific chemical proteomic substrate profiling

The chemical genetic system described above offers the first opportunity to discover ZDHHC/substrate networks *de novo* through chemical proteomics, by coupling metabolic labeling to enrichment and quantitative proteomics. Taking 18-Bz/ZDHHC20[Y181G] for our proof of concept, we first optimized selectivity for loading and transfer over 18-Bz/WT ZDHHC20, with respect to probe concentration and time. Loading of ZDHHC20[Y181G], but not WT ZDHHC20, increased with 18-Bz concentration, saturating at 20 μM 18-Bz; ZDHHC20[Y181G] loading and IFITM3 transfer reached a maximum plateau at 4-8 hours, with a similar trend observed for global labeling measured by in-gel fluorescence analysis of TAMRA-labeled lysates (Extended Data Fig. 4). We further confirmed that HEK293T cell proliferation and viability remained unperturbed by 18-Bz or 20-Bz treatment over at least 3 days under a range of culture conditions (Extended Data Fig. 5), and whilst mutant-selective 18-Bz labeling was clearly observed in media supplemented with 10% fetal bovine serum (FBS), stronger labeling intensity could be achieved at lower serum levels (Extended Data Fig. 6). Direct conversion of 18-Bz to 18-Bz-CoA in cells was confirmed by LC-MS/MS analysis of extracted metabolites (Extended Data Fig. 7). Furthermore, lipidomic analysis of cells treated with 18-cPr and 18-Bz revealed incorporation of each probe into the cellular pool of structural and storage lipids, including phosphatidylethanolamine (PE), phosphatidylcholine (PC) and triglyceride (TG) species, a process which requires *in situ* activation to the CoA ester (Extended Data Fig. 8A-C). These analyses also revealed that treatment with bumped lipids caused no significant perturbations across all major classes of endogenous lipids relative to vehicle or YnPal treated cells (Extended Data Fig. 8D).

**Figure 4.**
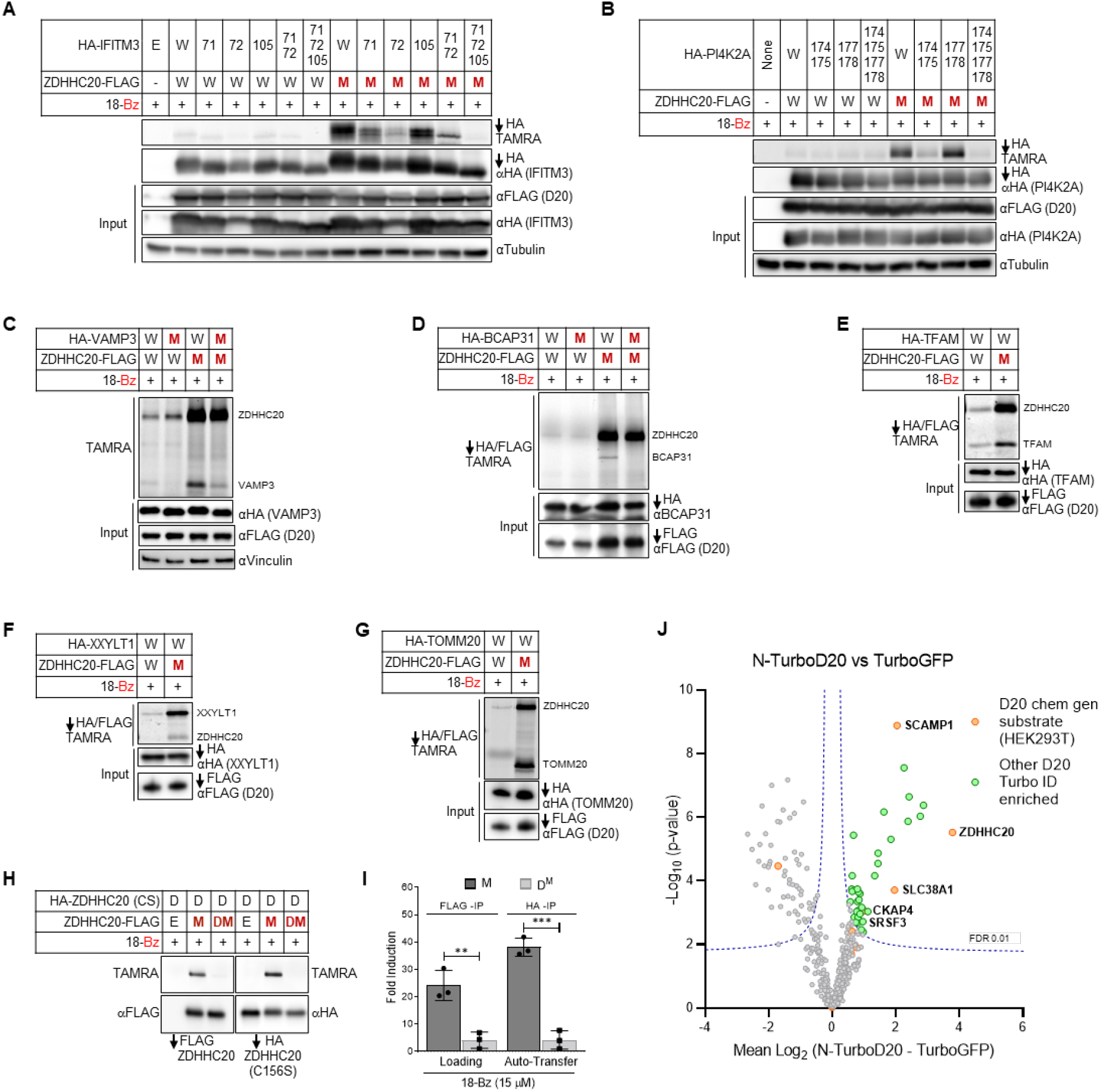
ZDHHC20 substrate and *S*-acylation site analysis. (**A**, **B**) ZDHHC20[Y181G] replicates reported selectivity profiles between specific cysteines on substrates IFITM3 and PI4K2A, with treatment of 15 μM 18-Bz for 4 h in HEK293T cells following immunoprecipitation, CuAAC with TAMRA azide, and analysis by in-gel fluorescence (n=3). (**C**, **D**) Validation of HA-VAMP3 and HA-BCAP31 *S*-acylation through fluorescence-based transfer assays by ZDHHC20[Y181G] with 15 μM 18-Bz treatment and *S*-acylation site mutants VAMP3[C76A] and BCAP31[C23A] in HEK293T cells (n=3). (**E**, **G**) Three novel *S*-acylated proteins, TFAM, XXYLT1, and TOMM20 validated through fluorescence-based transfer assays, demonstrating selective loading and transfer of 18-Bz in ZDHHC20[Y181G]-expressing HEK293T cells (n=3). (**H**, **I**) Auto-*S*-acylation *in trans* by ZDHHC20[Y181G] (M) of peripheral cysteines of catalytically dead ZDHHC20[C156S] (D) is shown using specific transfer of 18-Bz. Following treatment with 15 μM 18-Bz HEK293T cells were lysed and HA- and FLAG-tagged proteins were enriched separately by immunoprecipitation prior to CuAAC with TAMRA azide and analysis by in-gel fluorescence. (**I**) Average (n = 3) loading and transfer activity as percent maximal fluorescent/input ratio ± S.D; Prism 9.0 unpaired t-test statistical module was used to determine p-values (ns (not statistically significant), * P > 0.05, ** P ≤ 0.05, and *** P ≤ 0.01). (**J**) Example of detection of ZDHHC20 (D20) interactors through TurboID proximity-labeling; plot shows mean Log_2_ difference in protein group intensities between N-terminal TurboID-ZDHHC20 and TurboID-GFP clones; previously identified ZDHHC20 substrates in orange, significantly enriched proteins in green (Student’s unpaired T-test, S_0_ 0.1, FDR 0.01, n=4 per condition, see also Extended Data Fig. 12).

**Figure 5.**
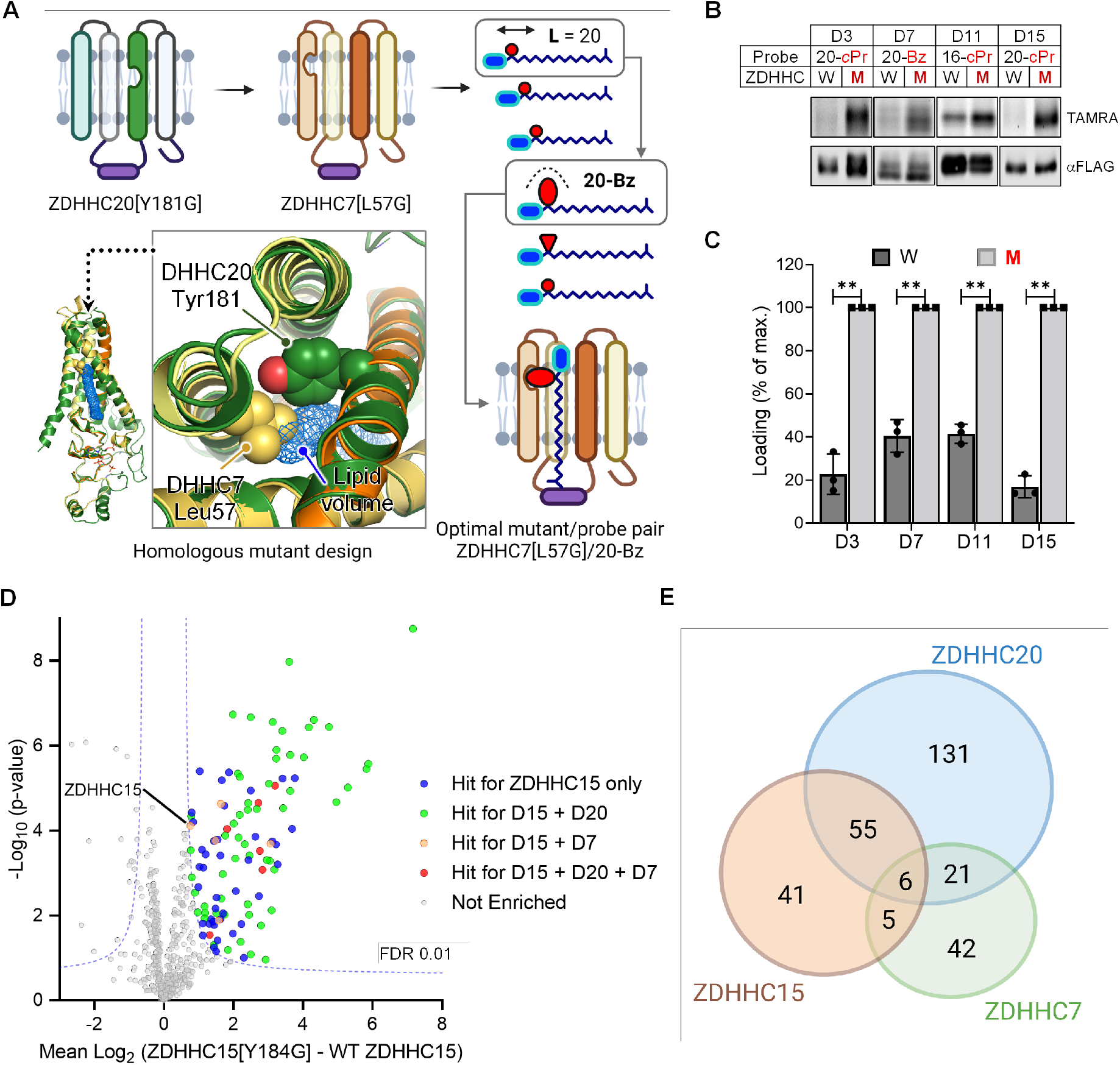
Extension of ZDDHC chemical genetics to ZDHHC3, 7, 11 and 15. (**A**) Structure-guided ZDHHC engineering exemplified for ZDHHC7 (see also Extended Data Figs. 13-16); ZDHHC7 homology model (yellow/orange) overlayed on experimental ZDHHC20 structure (dark green) identifies a potential hole-generating amino acid (Leu57) on an adjacent helix in the vicinity of ZDHHC20 Tyr181; lipid density (blue mesh), and length/size probe analysis identifies a mutant/probe pair (ZDHHC7[L57A]/20-Bz) with optimal activity and selectivity over wild type ZDHHC7. (**B**) Bump-hole analysis of N-FLAG-tagged WT ZDHHCs or mutant ZDHHCs ZDHHC3[I182G] (D3), ZDHHC7[L57G] (D7), ZDHHC11[M181A] (D11) and ZDHHC15[Y184G] (D15) in HEK293T cell-based loading assays using 15 µM corresponding optimized probe. (**C**) Average (n = 3) loading reported as a percent of maximal fluorescent:input ratio ± S.D.; p-values determined by Prism 9.0 unpaired t-test statistical module; ***, P ≤ 0.001 and ****, P ≤ 0.0001. (**D**) ZDHHC15 substrate discovery in HEK293T cells treated with 15 µM 20-*c*Pr in HEK293T cells using the OBH workflow; 107 chemical genetic ZDHHC15 substrates were identified (Student’s unpaired T-test, S_0_ 0.5, FDR 0.01, n=4 per condition); substrates unique or in common with parallel analyses for DHHC7 and DHHC20 in HEK293T cells are highlighted (see Extended Data Figs. 17 and 18). (**E**) Overlap of chemical genetic ZDHHC substrates identified in HEK293T cells. Of 301 total substrates, only 87 are shared by 2 or more family members, suggesting distinct substrate pools for each ZDHHC.

We next sought to discover substrates and *S*-acylation sites modified by ZDHHC20 *de novo* through quantitative mass spectrometry-based analysis of proteins labeled by 18-Bz in cells expressing either ZDHHC20[Y181G] or WT ZDHHC20^24^. We also aimed to determine specific sites *S*-acylated by ZDHHC20 simultaneously in the same samples, through on-bead thioester hydrolysis (OBH) and differential cysteine capping (Fig. 3A)^38^. HEK293T cells transfected with WT ZDHHC20 or ZDHHC20[Y181G] were treated with 15 μM 18-Bz for 8 h, and proteins recovered following lysis were subjected to CuAAC ligation with azido biotin capture reagent (AzB). Following CuAAC ligation, free (non-acylated) cysteines were blocked by N-ethylmaleimide (NEM) treatment, and labeled proteins enriched on neutravidin resin. Thioester linkages were then hydrolyzed on-bead by hydroxylamine treatment and these previously *S*-acylated cysteine residues capped with chloroacetamide (CAA). This strategy not only reduces contamination from non-specifically bound proteins, but also enables identification of sites *S*-acylated by ZDHHC20 in the same experiment marked by cysteine carbamidomethylation, following off-resin trypsin digestion and nanoLC-MS/MS analysis (Fig. 3A).

Label-free quantification (LFQ) revealed that 213 proteins were significantly enriched from HEK293T cells expressing ZDHHC20[Y181G] but not in ZDHHC20 WT. Furthermore, 99 potential *S*-acylation sites were identified (Fig. 3B-C and Supplementary Excel Tables 1, 2), including ZDHHC20 auto-*S*-acylation^35^ and 28 sites previously reported in targeted and global *S*-acylation studies, e.g. Cys11 and Cys15 of CD151 (Fig. 3C, Extended Data Fig. 9), consistent with detection of genuine sites of *S*-acylation^6, 39^. Known ZDHHC20 substrate IFITM3 was among the most significantly enriched proteins (>16-fold for ZDHHC20[Y181G] over WT) (Fig. 3B), demonstrating a level of sensitivity sufficient to identify ZDHHC20 substrates present at endogenous expression levels. We further extended chemical proteomic substrate identification to MDA-MB-231 and PANC1 cell lines, identifying 50 and 200 substrates, respectively, alongside 89 sites of modification (Extended Data Fig. 10, Supplementary Excel Table 1, 2), underscoring the versatility and adaptability of the system. Whilst 104 substrates were common to at least two out of three cell lines (HEK293, MDA-MB-231 and PANC1, Fig. 3D), we also identified unique sets in individual cell lines, which may indicate differences in substrate expression or context-specific regulation of *S*-acylation by ZDHHC20.

To explore localization and functional diversity across the first chemical genetic ZDHHC20 substrate maps we performed overrepresentation analyses using PANTHER, focusing on gene ontology (GO) terms for cellular compartments (slim) or biological processes. Substrates appearing in at least two cell lines were statistically overrepresented (FDR < 0.05) compared to genes from the reference human genome in compartments including the endomembrane system, ER, plasma membrane and intracellular vesicles, consistent with the reported cellular localization of ZDHHC20 (Fig. 3E)^18, 40, 41^. Analysis of biological processes compared to a reference database of known *S*-acylated proteins^6^ revealed enrichment of transport and glycosylation, and a profile distinct from the *S*-acylated proteome as a whole (Fig. 3F and Methods), consistent with a differentiated set of ZDHHC20 substrates.

### ZDHHC20[Y181G] conserves native acylation site specificity and reveals novel substrate modification sites

We next explored the biochemical conservation of the site-specificity of chemical genetic *S*-acylation compared to WT ZDHHC20-mediated *S*-acylation of established *S*-acylated substrates IFITM3 and PI4K2A by Ala scanning mutagenesis of known *S*-acylated cysteines. 18-Bz/ZDHHC20[Y181G] labeling showed specific *S*-acylation patterns exactly in line with those previously reported for IFITM3 and PI4K2A (Fig. 4A, B)^42, 43^. To validate novel *S*-acylation sites for ZDHHC20 substrates (Fig. 3B) we mutated the sole cysteines of VAMP3 and BCAP31 to Ala and confirmed *S*-acylation of VAMP3 at Cys76, and BCAP31 at Cys23 (Fig. 4C-D). Additionally, chemical proteomics analyses identified several putative novel ZDHHC20 substrates, TFAM, XXYLT1 and TOMM20, not previously reported as *S*-acylated proteins. 18-Bz/ZDHHC20[Y181G] labeling validated these proteins as *S*-acylated substrates for the first time, and furthermore linked their *S*-acylation specifically to ZDHHC20 (Fig. 4E-G). Interestingly, several ZDHHC20 sites (*e.g.* Cys263) were also identified, suggesting that ZDHHC20 may auto *S*-acylate *in trans* at non-catalytic sites. To test this hypothesis, we compared labeling of active versus catalytically inactive ZDHHC20 in HEK293T cells co-transfected with orthogonally HA-tagged inactive ZDHHC20[C156S] (D), and either empty vector (E), FLAG-tagged active (M), or inactive (D^M^) ZDHHC20[Y181G]. Immunoprecipitation using anti-FLAG or anti-HA beads showed that only active ZDHHC20[Y181G] is loaded with 18-Bz and can transfer probe to the HA-tagged inactive counterpart, confirming auto-*S*-acylation of ZDHHC20 *in trans* (Fig. 4H-I).

### Chemical genetics complements conventional approaches to ZDHHC substrate discovery

ZDHHC20 substrates identified through chemical genetics were next compared with conventional substrate identification approaches, including chemical proteomic analyses in ZDHHC20 knockout (KO) cell lines and interactome analyses through proximity biotinylation by TurboID-fused ZDHHC20^44^. We compared *S*-acylation of IFITM3 and metabolic labeling with YnPal against WT ZDHHC20 in two clonal ZDHHC20 KO HEK293T cell lines generated by CRIPSR/Cas9 (Extended Data Fig. 11). No changes were apparent in IFITM3 *S*-acylation between KO and WT cells (Extended Data Fig. 11B-C). Similarly, very few significant changes were observed in chemical proteomic analyses of *S*-acylated proteins using YnPal, with approximately equal numbers of enriched or depleted proteins (Extended Data Fig. 11D-F, Supplementary Excel Table 3). Quantitative interactome analyses of either N- or C-terminal ZDHHC20 TurboID fusions vs. TurboID-GFP in HEK293T cells (Extended Data Fig. 12) together identified only five proteins also identified as ZDHHC20 chemical genetic substrates (Fig. 4J, Extended Data Fig. 12D), consistent with the common observation that transferase substrates are not typically strong interactors^45, 46^. Taken together, these data suggest that ZDHHC chemical genetics offers a complementary approach to existing technologies by circumventing redundancy within ZDHHC substrate networks, whilst enhancing specificity and sensitivity toward *bona fide* substrates.

### Expansion of ZDHHC chemical genetics to multiple diverse family members

We next sought to apply chemical genetics to other members of the ZDHHC family. Models for the 4TM helices defining the lipid-binding pocket and catalytic domain were generated for each of the remaining 22 human ZDHHCs using a crystal structure of ZDHHC20 as a template, to identify suitable residues for mutagenesis and activity studies (Extended Data Fig. 13A)^3^. Guided by B-factors in the ZDHHC20 crystal structure, bulky residues buried in rigid hydrophobic pockets were prioritized above mutations on flexible structures such as loops since these are more likely to present suitable sites for steric complementation^47^ (Extended Data Fig. 13B) alongside alignment to ZDHHC20 Tyr181 (Extended Data Fig. 13C). We deprioritized potentially destabilizing mutations at the helix-bilayer boundary and cross-referenced models against AlphaFold structure predictions for the 4TM lipid binding core in the 6TM ZDHHCs 13 and 17, with root-mean-square deviations of 1.5 and 1.7 Å, respectively (Extended Data Fig. 14A-B)^48^. After identifying target residues for mutagenesis, we generated Ala, Gly or double mutant constructs for each ZDHHC and subjected them to the same two-stage screening strategy employed for ZDHHC20 (Fig. 5A).

For several ZDHHCs, Gly mutation resulted in dramatically decreased expression (Extended Data Fig. 15), which in some cases could be restored by switching from Gly to a structurally more conservative Ala mutation, or by adding strategically designed rescue mutations. For example, M181A in place of M181G improved ZDHHC11 expression, potentially by restoring backbone rigidity (Extended Data Fig. 15J-K). ZDHHC15[Y184G] was also only modestly expressed; the reported crystal structure of zebrafish ZDHHC15 showed Ser35 hydrogen bonding to Tyr184, suggesting that Y184G may expose the Ser35 hydroxyl to the lipid binding pocket. In support of this hypothesis, replacement of Ser35 with a less polar residue in a ZDHHC15[Y184G/S35C] double mutant improved expression (Extended Data Fig. 15O-P). In the length screen, we identified several probes displaying either selectivity or similar loading capacity for mutant versus WT ZDHHC; for example, 20-Ac was selectively loaded by ZDHHC3[I182G] and by ZDHHC15[Y184G/S35C], whilst 16-Ac was selectively loaded by ZDHHC11[M181A] (Extended Data Figs. 15C, 15K, 15P). Further refinement of probe selectivity for mutant over WT ZDHHC through bump screening at the optimal chain length led to the discovery of four new optimized mutant/probe pairs: ZDHHC3[I182G]/20-*c*Pr, ZDHHC7[L57G]/20-Bz, ZDHHC11[M181A]/16-*c*Pr and ZDHHC15[Y184G]/20-*c*Pr (Fig. 5A-C, Extended Data Fig. 16A-H), following optimization of expression to match WT ZDHHC. Taken together these data demonstrate that structure-guided design can expand chemical genetics across diverse ZDHHC family members, and we anticipate that future generations of optimal ZDHHC mutant/probe pairs can be established by combining refined modeling approaches with wider ranging mutant screens and additional bumped lipid designs.

### Chemical genetics enables comparative ZDHHC substrate profiling

The identification of diverse ZDHHC mutant/probe pairs offers the opportunity to undertake comparative substrate profiling between ZDHHCs for the first time. Chemical proteomic analyses of ZDHHC7[L57G]/20-Bz and ZDHHC15[Y184G]/20-cPr in HEK293T cells identified 74 and 107 substrates, respectively (Extended Data Fig. 17, Supplementary Excel Tables 5 and 6), alongside 20 sites of modification across 13 proteins (Supplementary Excel Table 2). Similarly, ZDHHC15 profiling in PANC1 cells rendered 91 substrates, 41 of which in common with those identified in HEK293T cells (Extended Data Fig. 17C-D, Supplementary Excel Table 5).

Among the 301 unique substrates identified across the chemical proteomic profiles, we observed common and distinct substrates between ZDHHCs 7, 15 and 20 consistent with a degree of redundancy across the DHHC family^29, 49^ (Fig. 5D and 5F, Extended Data Fig. 18). However, common substrates were in the minority, with 55 (24%) shared substrates between closely related ZDHHC15 and ZDHHC20 (48% sequence identity), and only six (PTPN1, RHBDD2, SCAMP2, SLC7A1, TMEM161A, HMOX2) shared by all three *S*-acyltransferases. These data suggest that chemical genetics combined with chemical proteomics provides the first direct approach to evaluate and compare substrate scope between specific ZDHHCs in intact cells.

## Discussion

Chemical genetics opens a new ZDHHC-specific window on the expansive *S*-acylation network, with the possibility to enhance detection of substrates and *S*-acylation sites with low abundance or stoichiometry, whilst simultaneously linking them to a cognate ZDHHC by circumventing ZDHHC redundancy. For example, HEK293T cells express all 23 human ZDHHCs to a measurable extent^50^, and against this background IFITM3 *S*-acylation is essentially unaffected by ZDHHC20 knockout, consistent with previous studies suggesting that IFITM3 may be *S*-acylated by ZDHHCs 3, 7, 15, and 20^21^. Indeed, a combination of traditional ZDHHC20 substrate identification strategies (knockout, overexpression, or proximity labeling with N- or C-terminal TurboID fusions) identified few significant hits and failed to identify known substrates at endogenous abundance. In contrast, chemical genetic analyses readily identified endogenous IFITM3 as a high confidence ZDHHC20 substrate.

We envisage further extending chemical genetic systems across the *S*-acyltransferase family to generate comprehensive contextual ZDHHC-specific substrate maps analogous to kinase-specific phosphorylation datasets, enabling elucidation of substrates and sites common and unique between ZDHHC isozymes in varied cell and tissue types. The present study illustrates the potential of this approach through identification of diverse partially overlapping substrate sets which exhibit a narrower spectrum of functional annotation than the wider *S*-acylated proteome. Validation of novel and established ZDHHC20 substrates alongside high fidelity recapitulation of known *S*-acylation site stoichiometry demonstrates that these datasets encompass *bona fide* ZDHHC-specific substrates, alongside a rich set of putative substrates for future validation. Chemical genetics also offers a unique approach for resolving ZDHHC isoform-dependent *S*-acylation at the level of specific PTM sites, whilst limiting or eliminating probe distribution into non ZDHHC-dependent pathways (e.g. membrane lipid biosynthesis, auto-*S*-acylation, and *O*- and *N*-linked acylation), an unavoidable liability of generic lipid analogues such as YnPal^38, 51^. Lipidomic analyses suggest that bumped probes are not extensively processed into membrane lipid pools, and do not alter endogenous lipid biosynthetic pathways; however, further probe optimization may be required to enhance activation and increase delivery via mutant ZDHHCs, particularly at higher serum levels.

Whilst we have demonstrated a systematic approach to establish effective chemical genetic systems for five ZDHHCs (3, 7, 11, 15 and 20), including strategies for rescue mutations, the formidable challenges of ZDHHC crystallization present a bottleneck for structure-guided design of complemented ZDHHC/probe combinations^3, 34^. Model refinement through machine learning ^48^, deeper mutational analysis, and design of additional bumped lipid probes offer expansive opportunities for optimization of progressively more efficient chemical genetic systems for each of the human ZDHHCs. Next generation designs may overcome some of the limitations of the current implementation whilst enabling new applications, for example, analysis in knock-in lines expressing mutant ZDHHCs at wild type levels or cell-type specific analysis of ZDHHC activity in organoid or animal models^52^. Compatibility of this system with cellular acyl protein thioesterase (APT) activity, which acts to reverse *S*-acylation, should also be investigated. Future systems bearing additional orthogonal tags may permit multiplexed analysis of two or more ZDHHCs in the same cell to reveal co-regulation and ZDHHC cascades, as previously disclosed for ZDHHC6 and ZDHHC16^53^. Previous studies have associated dysregulation of ZDHHC activity with diverse pathologies including cancer, inflammation and neurodegeneration, and we envisage applications of chemical genetics for drug target validation and discovery in ZDHHC-associated disease models, and in diverse disease-causing eukaryotes, for example parasites or fungi. Chemical proteomics also offers an ideal platform to analyze in-family selectivity for future small molecule ZDHHC inhibitors; conversely, it may prove possible to adapt bumped probes into chemical genetic inhibitors, offering a general solution to the current lack of specific ZDHHC inhibitors for functional studies.

## Figures

## Methods

### Cell Culture and Compound Preparation

HEK293T, HEK293-FT, MDA-MB-231 and PANC1 cell lines were cultured in Dulbecco’s Modified Eagle Medium (DMEM) supplemented with GlutaMAX^TM^, 10% v/v FBS, 100 U/mL penicillin and 0.1 mg/mL streptomycin in a 37^°^C, 5% CO_2_ incubator. Puromycin dihydrochloride (Life Technologies) and Blasticidin S hydrochloride (Cambridge Bioscience) stock solutions were prepared by diluting antibiotics with milli-Q H_2_O to a concentration of 10 mg/mL and stored at -20^°^C. Synthetic lipid-based loading and transfer probes and alkyne palmitic acid (2B Scientific BCAL-015-25), Palmostatin B (Sigma-Aldrich), TAMRA or biotin PEG_3_ azide (Sigma-Aldrich, 760757 or 762024), tris[(1-benzyl-1*H*-1,2,3-triazol-4-yl)methyl]amine (TBTA, Sigma-Aldrich, 678937) and palmitoyl coenzyme A (≥ 90%, Sigma-Aldrich, P9716) were dissolved in DMSO and stored at -20^°^C. Sealed ampules containing a 0.5 M aqueous solution of tris (2-carboxyethyl) phosphine (TCEP) and TCEP HCl (C4706) were purchased from Sigma-Aldrich and n-Dodecyl-β-D-maltopyranoside (DDM) was purchased from Generon (D310LA). TCEP HCl was prepared fresh as 50 mM stock in Milli-Q H_2_O, DDM was prepared as a 10% stock solution in Milli-Q H_2_O and stored at -20^°^C and cOmplete^TM^, EDTA-Free Protease Inhibitor Cocktail tablets were used according to the manufacturer’s instructions (Sigma-Aldrich, 11873580001).

### Antibodies and Western Analysis

Mouse-derived monoclonal antibodies for FLAG^®^ M2 (F1804), HA-epitope (HA-7, H3663) and α-tubulin (T5168) and rabbit-derived polyclonal antibodies for ZDHHC20 (Atlas Antibodies, HPA014702), BCAP31 (Atlas Antibodies, HPA003906) and V5-epitope (SAB1306079) were purchased from Sigma-Aldrich. Mouse monoclonal anti-GFP (GF28R) antibody was purchased from Generon LTD and rabbit-derived polyclonal antibodies against vinculin (42H89L44) and calnexin (ab22595) were purchased from Thermo Fisher Scientific and Abcam, respectively. Secondary antibodies were HRP-conjugated, polyclonal goat-derived antibodies against mouse and rabbit (Dako, Agilent Technologies) and IRDye 800CW goat anti-mouse (ab216772). Western blot analysis was accomplished through SDS-PAGE of cell lysates or affinity-resin eluates in 1X laemmli loading-buffer (Bio-Rad) containing β-mercaptoethanol (1/10 in 4X buffer) and transfer of protein onto PVDF or nitrocellulose using the Trans-Blot Turbo System (Bio-Rad). Secondary HRP-conjugates were visualized after addition of ECL Prime Western Blotting Substrate and chemiluminescent detection in an Amersham Imager 680 or fluorescence detection using a LICOR Odyssey CLx. Quantitation of western blot protein intensity was performed by densitometry using ImageJ 1.50c or Image Studio Lite and data were plotted using Prism 9.0.

### ZDHHC Structural Modelling

Human ZDHHC family protein sequences were aligned using the ‘Create Alignment’ module of CLC Sequence Viewer 7. Regions of sequence similarity that also overlap with ZDHHC20 transmembrane helices (TMs) 1 – 4 and the DHHC-containing cysteine-rich domain were identified and selected for homology modelling. In order to generate homology models for ZDHHCs 1 – 19 and 21 – 24, selected sequences were individually submitted to the **P**rotein **H**omology/analog**Y R**ecognition **E**ngine V **2**.0 (Phyre^2^) using the ‘Normal’ modelling mode ^54^. In order to identify putative bump-hole mutations, homology models were structurally aligned to the ZDHHC20-2BP crystal structure (PDB entry **6BML**) using MacPyMOL: PyMOL V1.5.0.4. Residues located on TM3 and spatially overlapping with or proximal to ZDHHC20-Y181 were prioritized for bump-hole analysis; however, several ZDHHC-models did not present residues meeting these criteria. In this case, strict rules were adopted including: 1) Selection of ZDHHC20-Y181 or 2) -F65 proximal residues located on TMs 2 and 3 overlapping with ZDHHC20 residues having B-factor values < 100 and, 3) when rules 1 and 2 fail, residues on TM1 proximal to the ω-position of the 2BP fatty-acid chain and with lowest ZDHHC20 B-factor value were selected for bump-hole screening. Residues presented by TM1 were given least priority as their side chains have access to the lipid-binding pocket and the bilayer. Potentially, mutations on this helix could generate a hole in the pocket, leading to structural instability or loss of lipid-probe binding-affinity.

### Molecular Biology and Cloning

#### Plasmids and Subcloning

For a complete list of vectors used in this study, refer to Supplementary Table 2. The preparation of new plasmids generated for this study will be described herein. Human C-terminally Myc-FLAG-tagged ZDHHC20 (C-FLAG-D20) was purchased from Origene Technologies, catalogue number MR205665. Plasmids for expression of N-terminally 3xFLAG-tagged human ZDHHCs 1 – 24 (N-FLAG-DX, X = 1 – 24), HA-tagged GCP16 (GOLGA7, ZDHHC9 cofactor) and empty pEF-1α vector were a generous gift from Y. Ohno (Hokkaido University). The C-FLAG-pcDNA3 expression vector (Addgene 20011) was also used as a negative control for FLAG-tagged ZDHHC expression. C-terminally Myc-HA-tagged ZDHHC20 (C-HA-D20) PCR fragment was subcloned into the PmeI and AsiSI linearized C-FLAG-D20 vector using the NEBuilder HiFi Assembly Kit. C-HA-D20 fragment, with HA-tag sequence spacer (**bold**), was generated by PCR using Phusion DNA Polymerase and the primer set in Supplementary Table 3. C-FLAG-D20 was subcloned into the pLVX-TetOne-Puro (Clontech) and in-house attb vectors, expressing mouse ZDHHC20 and containing a blasticidin resistance marker (attb-ZDHHC20-BSDr), using the same strategy. Empty pLVX-TetOne-Puro and attb-ZDHHC20-BSDr were linearized using EcoRI and BamHI and XhoI and MfeI, respectively, and human C-FLAG-D20 fragments generated with primer sets indicated in Supplementary Table 3. V5-tagged TurboID (promiscuous BirA mutant, V5-Turbo-NES-pCDNA3, Addgene 107169) and C-FLAG-D20 or EGFP (pEGFP-N1-FLAG, Addgene 60360) chimeras were also subcloned into the XhoI and MfeI linearized attb-BSDr vector using the NEBuilder HiFi Assembly Kit; See Supplementary Table 4 for primer sets. Plasmids for the expression of PI4K2A (pDONR223-PI4K2A Addgene 23503), TOMM20 (mCherry-TOMM20-N-10 Addgene 55146) and TFAM (pcDNA3-TFAM-mCLOVER Addgene 129574) were subcloned into the EcoRI and XhoI linearised N-HA-IFITM3 vector using the NEBuilder HiFi Assembly Kit; see Supplementary Table 3 for primer sets used to create N-HA tagged plasmids. pcDNA3.1 XXYLT1-HA was kindly provided by Dr Hans Bakker. All new plasmids were sent to GATC Biotech for Sanger sequencing to confirm sequences of entire inserts and junctions between backbone and insert and backbone and PCR fragment. Primers utilized to produce PCR fragments were purchased from Sigma-Aldrich.

#### Molecular Cloning

Lentiviral plasmid pLVX-C-FLAG-D20, packaging and viral envelope plasmids pCMV-Delta-8.2 (Addgene 12263) and pCMV-VSV-G (Addgene 8454) and HEK293-FT cells were used to prepare lentiviral particles according to the instructions in the Lenti-X^TM^ Lentiviral Expression System Manual (Clontech). Doxycycline-inducible ZDHHC20 expressing HEK293T cells were prepared by transduction of 2.5 X 10^5^ low-passage cells with lentivirus, followed by puromycin selection at 1.0 μg/mL for 1 week. After 1 week, cells were cultured in normal media containing 0.5 μg/mL puromycin and, after 4 or more passages, sent to the Flow Cytometry Science Technical Platform for single-cell sorting into 96-well plates. Briefly, single cells were sorted on the Beckman Coulter MoFlo XDP, using the 488 nm forward scatter (FSC) LASER signal to trigger events. Cells were identified and separated from debris using side scatter (SSC) height vs. FSC height. Doublets were then removed using SSC height vs SSC width (Extended Data Figure 19A-B). Single cells were sorted into 96-well plates using the Single Cell sort mask with a drop envelope of 0.5. Clones were expanded into 12-well plates and induced with 2 μg/mL doxycycline for 2 days before anti-FLAG western blot screening. Jump-in TurboID cell lines were prepared by co-transfecting HEK293T cells with 1.0 μg and 1.5 μg of attb-BSDr containing the gene of interest and pCMV-Int (ΦC31-integrase, Addgene 18935) plasmids respectively, followed by selection with 10 μg/mL blasticidin for 1 week. For transfection conditions, see **Cellular and Biochemical Analysis:** *Cellular ZDHHC-autoacylation and substrate-transfer analysis* section. After selection, single cells were sorted into 96-well plates, as described above, expanded into 12-well plates and then screened by immunoblot with antibodies against GFP, V5 and FLAG epitopes.

#### Subcloning BCAP31 into Mammalian Expression Vector

After extracting total RNA from HEK293T cells using the GenElute Total RNA Purification Kit, 500 ng of total RNA was used to amplify BCAP31 (isoform 1, P51572) cDNA using the SuperScript III One-Step RT-PCR System with Platinum Taq DNA Polymerase and the following primer set: forward primer ATG AGT CTG CAG TGG ACT GCA GTT G and reverse primer TTA CTC TTC CTT CTT GTC CAT G. BCAP31 cDNA was purified by agarose gel electrophoresis, extracted and shuttled into a TOPO-TA vector. After blue-white screening, white colonies were selected, amplified in the presence of ampicillin and then harvested to prepare DNA minipreps. Miniprepped DNA was digested with KpnI and SmaI and analyzed by agarose gel electrophoresis to confirm the presence of BCAP31; all inserts were oriented in the reverse-sense and Sanger sequencing verified the sequence of human BCAP31, isoform 1. To generate an HA-tagged BCAP31 construct in a mammalian expression vector, BCAP31 PCR fragment was subcloned into SalI and BalI linearized pcDNA3.1(+)-HA-IFITM3 vector using the NEBuilder HiFi Assembly kit. PCR primers used to generate the HA-BCAP31 PCR fragment are listed in Supplementary Table 3.

#### Mutagenesis

All mutagenesis reactions were carried out using the QuikChange II Site-Directed or Lightning MultiSite-Directed Mutagenesis Kits (Agilent) according to the manufacturer’s instructions. See Supplementary Tables 5 and 6 for a list of mutations and mutagenic primer sets. All plasmids were sent to GATC Biotech or Genomics Equipment Park (The Francis Crick Institute) for Sanger sequencing to confirm mutations. Mutagenic primers were purchased from Sigma-Aldrich.

### Cellular and Biochemical Analysis

#### Cellular ZDHHC-autoacylation and substrate-transfer analysis

FLAG-tagged ZDHHC was singly or co-transfected with affinity-tagged substrate (FLAG or HA epitopes) in HEK293T cells. For a single, reverse-transfection mixture, 1 – 2 μg of ZDHHC plasmid, alone (autoacylation) or in combination with 0.5 μg substrate (transfer) plasmid, was mixed with 3 volumes of Fugene HD (3:1 Fugene (μL)/Plasmid (μg)) in 100 μL of Opti-MEM. After 15 min, the transfection mixtures were added to 1 – 2 X10^6^ cells in 900 μL of culture media and cells incubated at 37°C o/n. Cells were treated with DMSO or probe in 10% FBS (YnPal) or 0.5% FBS (bumped probes) for 4 h at 37^°^C, after which they were dislodged by manual pipetting, pelleted at 200 x g for 5 min and the media discarded. Cells were washed 2X ice cold PBS and pelleted each time to remove the supernatant. Cells were lysed with 0.5 mL DDM lysis buffer: 50 mM Tris-HCl, pH 7.5, 150 mM NaCl, 10% glycerol, 1% DDM, 0.5 mM TCEP, 10 μM Palmostatin B, 1X protease inhibitor tablet, 2 mM MgCl_2_ and 0.05 U/μL Benzonase. After shaking for 10 min at RT, lysates were clarified at 17,00 x g for 10 min at 4°C. The clarified lysates were treated with the appropriate affinity resins; anti-FLAG M2 (Sigma-Aldrich, M8823) and/or Pierce anti-HA (Life Technologies, 88836) magnetic beads, or diluted with 4X laemmli buffer for analysis. FLAG and/or HA-tagged proteins were immunoprecipitated for 2 h at RT or o/n at 4^°^C, then washed with wash buffer (1% NP-40 in PBS). For dye incorporation, 15 μL of click-mix, composed of 100 μM TAMRA azide, 1 mM CuSO_4_, 1 mM TCEP and 100 μM TBTA in wash buffer, was added to the beads. Click-reactions were allowed to shake at RT for 1 h before the beads were washed again with wash buffer, treated with 1X Laemmli loading-buffer and then eluates subjected to SDS-PAGE. For samples used to demonstrate thioester dependent labelling, prior to 1X laemmli addition, beads were incubated with 0.8 M NH_2_OH in PBS pH 7.4 for 1 hour at RT then diluted with 4X laemmli buffer. After protein separation, TAMRA-labelled proteins were visualized using a Typhoon FLA 9500 instrument. The Typhoon resolution was set to 25 μm and the PMT value varied from 500 – 1000 depending on signal intensity. Finally, protein was transferred to PVDF/nitrocellulose and analysed via western analysis to reveal ZDHHC and substrate input. For loading and transfer, the fluorescent signals were normalized against input as determined by western blot analysis.

#### ZDHHC20 Enzyme Kinetics: Mutant and Probe Analysis

##### Protein Purification

For each FLAG-ZDHHC20 construct, two dishes of HEK293T cells were prepared and transfected with calcium phosphate transfection mix. To a 10 cm tissue culture dish was added 6 X 10^6^ HEK293T cells in 8 mL DMEM culture media. Cells were allowed to settle o/n in a 37°C incubator. To prepare the calcium phosphate transfection mix, 10 μg of C-FLAG-D20 construct and water was added to a 1.5 mL Eppendorf tube to a volume of 436 μL. To this was added 64 μL of 2 M CaCl_2_ to give a final volume of 500 μL. This solution was slowly added dropwise and with continuous bubbling to 500 μL 2X HBSS (273.8 mM NaCl, 9.4 mM KCl, 1.5 mM Na_2_HPO_4_-7H_2_O, 15 mM glucose, 42 mM HEPES (free acid) pH 7.05 in MilliQ-H_2_O, filter sterilized) in a sterile 30 mL polystyrene tube. This solution incubated at RT for 5 min before use. To a 10 cm plate of HEK293T cells was added 1 mL of the DNA mixture. The mix was added dropwise and evenly across the media surface to ensure maximal cell coverage with precipitated DNA. The tissue culture dish was returned to the 37^°^C incubator and left to incubate for 72 h before harvesting cells for protein purification.

After 72 h, cells were dislodged manually using an automatic pipette, collected in a 50 mL falcon tube and pelleted at 200 x *g* for 5 min. The media was decanted, and cells washed 3X with cold PBS. The cell pellet was dislodged and lysed with 5 mL 2% DDM buffer (50 mM Tris-HCl, pH 7.5, 150 mM NaCl, 5% glycerol, 2% DDM, 0.5 mM TCEP, 10 μM palmostatin B and 1X protease inhibitor tablet), vortexed thoroughly and allowed to incubate at 4^°^C with constant rotation. After 4 h, lysate was centrifuged for 20 min at 20,000 x *g* and 4^°^C. 1 mL anti-FLAG M2 Affinity Gel suspension was equilibrated in 5 mL 1% DDM buffer before use. The clarified lysate was diluted with 1 volume (∼ 5 mL) of the described buffer containing no DDM, to give a final concentration of 1% DDM, before being added to cold resin and allowed to mix o/n at 4^°^C. After incubation, the lysate-resin mix was added to an Econo-Pac gravity-flow chromatography column (Bio-Rad); the lysate container was rinsed thoroughly with 1% DDM buffer to transfer all residual resin to the column. The resin was washed sequentially with ice-cold buffer W1 (50 mM Tris-HCl pH 7.5, 150 mM NaCl, 0.2% DDM, 2 mM TCEP and 1X protease inhibitors), W2 (25 mM HEPES pH 7.5, 500 mM NaCl, 0.2% DDM, 2 mM TCEP and 1X protease inhibitors) and W3 (W2 with 25 mM NaCl). After washing, FLAG-ZDHHC20 was eluted with 5 X 0.5 mL portions of buffer W3 containing 0.25 mg/mL 3X FLAG-peptide (MDYKDHDGDYKDHDIDYKDDDDK, Crick Peptide Chemistry STP stock peptide). FLAG-ZDHHC20 containing fractions were pooled and buffer exchanged with W3 simultaneously with concentration using an Amicon Ultra 15 mL Spin-Column with 50 kDa cut-off.

#### Enzyme-coupled ZDHHC20-autoacylation assay

Autoacylation reactions were carried out as previously described with a few exceptions^3^. Reactions were prepared in a Corning 96-well black half-area plate. In one well was prepared a pre-start mix with the indicated concentration of fatty acid-or probe-CoA, 2 mM α-ketoglutaric acid, 0.25 mM NAD (β-nicotinamide adenine dinucleotide, oxidized) and 0.2 mM thiamine pyrophosphate in 25 μL reaction buffer (25 mM MES, pH 6.8, 50 mM NaCl, 1 mM DTT, 1 mM EDTA, 0.2 mM DDM and 1X protease inhibitors). The reaction was started by addition of 25 μL ZDHHC master mix containing 20 nM ZDHHC20 and 2 μL of α-ketoglutarate dehydrogenase (KDH, Sigma-Aldrich, K1502). In the first reaction step, ZDHHC20 autoacylation leads to production of free CoA-SH, which is then converted to NADH in the next step by KDH. NADH production was monitored using the fluorescence module (Ex. 360 nm/Em. 465 nm) of the Tecan Spark Multimode Microplate Reader. Note that 18-Bz-CoA exhibited weak, but measurable, activity in KDH reactions without ZDHHC20 enzyme. Therefore, all rates from reactions with ZDHHC20 and 18-Bz-CoA were adjusted by subtraction of the ZDHHC20-independent rates from the corresponding total rates of reactions with ZDHHC20 (Extended Data Fig. 3G).

#### Incucyte growth assays

Doxycycline (dox)-inducible FLAG-tagged WT and ZDHHC20[Y181G] expressing HEK293T cells were maintained in media supplemented with 2 μg/mL dox for 48 h. Following this, cells were seeded at a density of 5,000 cells/well into a 96-well plate. Cells were maintained at 2 μg/mL dox and treated in triplicate with either DMSO or 15 μM 18-Bz or 20-Bz at four different FBS concentrations: 0.5, 2, 5 and 10%. Cell confluence was measured in real-time using the Incucyte FLS Imaging System for 70.5 h. The cell confluence for each group was normalized to maximum value and plotted as the average (n = 3) % confluence ± S.D. using Prism 9.0.

## Metabolomics – see Supplementary Methods

### Quantitative Mass Spectrometry-Based Proteomics

#### On-bead hydrolysis proteomics

Cells were concurrently plated and transfected with 1 µg/mL of the appropriate ZDHHC construct using 3 µL of Fugene per 1 µg of DNA in a 10 cm dish. After 6 h the media was refreshed and cells were incubated overnight. Cells were treated with 15 µM of the appropriate probe in 0.5% FBS media for 8 h. Cells were then washed 3X with ice-cold PBS then lysed in SDS lysis buffer (50 mM HEPES pH 7.4, 0.5% NP-40, 0.25% SDS, 10 mM NaCl, 2 mM MgCl_2_, EDTA free cOmplete protease inhibitor (Roche) and 0.05 U/μl benzonase). Lysates were adjusted to 2 mg/mL, using ∼2 mg per condition, then subjected to a click reaction using biotin-PEG3-azide and small portion taken to be clicked with TAMRA azide, for analysis by western blot and in-gel fluorescence. The click reagent mixture was prepared by mixing 10 µL of 10 mM biotin-PEG3-azide, 20 uL of 50 mM CuSO_4_, 20 µL of 50 mM TCEP and 10 µL of 10 mM TBTA, 64 µL was added per 1 mL of lysate and the samples incubated for 1 h at RT with agitation. The reaction was quenched with 5 mM EDTA followed by chloroform/methanol precipitation.

For those samples where no site ID was performed protein samples were suspended in 50 mM HEPES pH 7.4 containing 1% SDS then precipitated with chloroform/methanol. For experiments including site identification of lipidation proteins were solubilised in 1 mL 50 mM triethanolamine pH 7.5, 4% SDS, 5 mM EDTA. TCEP (10 mM) was added to samples and incubated for 20 min at RT with agitation. To these solutions 25 mM *N*-ethylmaleimide (NEM, from 1 M stock in EtOH) was added and incubated for 2 h at RT with agitation. Samples were then precipitated with chloroform/methanol.

All protein samples were then suspended in 50 mM HEPES pH 7.4 containing 2% SDS then diluted to 0.2% SDS using 50 mM HEPES pH 7.4. Labelled proteins were enriched using NeutrAvidin agarose beads (Pierce) using 40 μL (50% slurry) per condition. The beads were equilibrated by washing in 0.2% SDS in 50 mM HEPES pH 7.4 prior to incubation with the samples for 3 h at RT with agitation. Beads were then washed twice with 0.2% SDS in 50 mM HEPES pH 7.4 and four times with 50 mM HEPES pH 7.4. Beads were then suspended in 20 µL of 50 mM triethanolamine pH 7.5, 4 mM EDTA and 0.5% ProteaseMAX (Promega). To this was added 10 µL of the NH_2_OH cleavage solution (100 µL 8.16 M NH_2_OH in 50 mM TEA pH 8.0, 30 µL 500 mM triethanolamine pH 7.5, 2.6 µL 500 mM EDTA, 197.4 µL water) giving a final NH_2_OH concentration of 0.82 M, and the samples incubated at RT for 2 h with agitation. Following this, 100 µL of 50 mM HEPES pH 8.0 containing 5 mM TCEP was added and the beads pelleted and 120 µL of supernatant taken. 20 mM chloroacetamide was added to the supernatant and they were incubated at RT for 15 min. Samples were then diluted with 400 µL 50 mM HEPES pH 8.0 and digested with 0.3 µg of trypsin (Promega) o/n at 37°C. Samples were acidified with 0.5% (v/v) trifluoroacetic acid (TFA), flash frozen and lyophilised. Samples were dissolved in water containing 0.5% TFA and loaded onto stage tips containing three SDB-XC poly(styrenedivinyl-benzene) copolymer discs (Merck). The stage tipping procedure was carried out as described by Rappsilber *et al*. ^55^ Peptide samples were eluted in 55% acetonitrile in water and the solvent removed by incubation in an Eppendorf Concentrator plus at 45°C.

Samples in which site ID was not performed were dissolved in water containing 0.1% TFA, ready for LC-MS/MS analysis. For samples where site ID was performed, prior to LC-MS/MS analysis samples were 3X fractionation using stage tips loaded with 3 layers SDB-RPS discs (3M^TM^ EmporeTM). Peptides were loaded on the solid phase in 0.5% TFA, and subsequently washed 3X with 60 µL 0.2% TFA. Peptides were eluted using 3 elution buffers (60 µL); buffer 1 (100 mM ammonium formate, 40% (v/v) MeCN, 0.5% (v/v) formic acid), buffer 2 (150 mM ammonium formate, 60% (v/v) MeCN, 0.5% (v/v) formic acid), buffer 3 (5% (v/v) ammonium hydroxide, 80% (v/v) MeCN). Samples were dried in an Eppendorf Concentrator plus at 45°C and then dissolved in water containing 0.1% TFA, ready for LC-MS/MS analysis. Peptides were analyzed on a Q-Exactive mass spectrometer (Thermo Fisher) coupled to an Ultimate3000 LC (Thermo Fisher) using an Easy Spray Nano-source. The instrument was operated in data dependent acquisition mode selecting the 10 most intense precursor ions for fragmentation.

#### YnPal KO Proteomics

6 x 10^6^ cells from HEK293T, HEK293T ZDHHC20 CRISPR knockdown cell line, and two HEK293T ZDHHC20 CRISPR knockout clonal lines were seeded in triplicate into 10 cm dishes and left o/n at 37°C. Cells were then treated with 15 μM YnPal or 15 μM palmitic acid and incubated for 8 h at 37°C. Cells were then washed 3X with PBS then lysed in SDS lysis buffer (50 mM HEPES pH 7.4, 0.5% NP-40, 0.25% SDS, 10 mM NaCl, 2 mM MgCl_2_, EDTA free cOmplete protease inhibitor (Roche) and 0.05 U/μl benzonase). Lysates were adjusted to 2 mg/mL, using 2 mg per condition, then subjected to a click reaction using biotin-PEG3-azide as described above. The reaction was quenched with 5 mM EDTA followed by chloroform/methanol precipitation. Protein pellets were washed and sonicated 2X with 1 mL MeOH.

All samples were then dissolved in 1% SDS in 50 mM PBSA pH 7.4 then diluted to 0.2% SDS with 50 mM PBSA pH 7.4. Biotinylated proteins were then enriched on a 1:1 mixture of dimethylated NeutrAvidin agarose beads (Pierce) ^56^ and control agarose beads (Pierce), which had been prewashed 2X 0.2% SDS PBSA pH 7.4, for 3 h at room temp. Beads were washed 3X 1 mL 0.2% SDS in PBSA followed by 2X 1 mL 50 mM HEPES pH 7.4 and finally 1X 1 mL 50 mM HEPES pH 8.0. Beads were then suspended in 50 µL 50 mM HEPES pH 8.0 with 400 ng of LysC (Promega) for 2 h at 37°C with agitation. The supernatant was removed and reduced with 5 mM TCEP and alkylated using 15 mM chloroacetamide for 15 min then digested with 100 ng of trypsin o/n at 37°C (Promega). Samples were acidified with 0.5% (v/v) trifluoroacetic acid (TFA) and the solvent removed in an Eppendorf Concentrator plus at 45°C. Samples were dissolved in water containing 0.5% TFA and after stage tipping using Oasis HLB μElution Plate 30 μm following the manufacturers procedure, and elutions were dried in an Eppendorf Concentrator plus at 45°C. Samples were dissolved in 2% MeCN, 97.9% water containing 0.1% TFA ready for LC-MS/MS analysis. Peptides were analyzed on a Q-Exactive mass spectrometer (Thermo Fisher) coupled to an Ultimate3000 LC (Thermo Fisher) using an Easy Spray Nano-source. The instrument was operated in data dependent acquisition mode selecting the 10 most intense precursor ions for fragmentation.

#### Proteomics Searches and Data Analysis proteomics OBH and YnPal

RAW files were uploaded into MaxQuant (version 1.6.7.0) and searched against Uniprot curated human proteome (As of 2019) using the built-in Andromeda search engine. Cysteine carbamidomethylation was selected as a fixed modification and methionine oxidation and acetylation of protein N terminus as variable modifications. For site ID experiments cysteine NEM modification and carbamidomethylation were selected as variable modifications. Trypsin/P was set as the digestion enzyme, up to two missed cleavages were allowed and a false discovery rate of 0.01 was set for peptides, proteins and sites with match between runs selected. Data was quantified using LFQ with a minimum ratio count = 2.

Data analysis was performed using Perseus (version 1.6.2.1). MaxQuant proteingroups.txt output files were uploaded and filtered against contaminants, reverse and proteins identified by site and a base 2 logarithm was applied to all LFQ intensities. For OBH, datasets were filtered to contain ≥3 valid values in the positive (mutant) condition. Missing values were imputed from a normal distribution (width = 0.3, downshift = 1.8). For YnPal samples, data was filtered for valid values in at least 2/3 of each condition for all analyses except when comparing against palmitic acid, where only 2/3 YnPal treated samples were considered and then missing values were imputed from a normal distribution (width = 0.3, downshift = 1.8) (Extended Data Fig. 11G). Within the YnPal datasets the data was normalized by subtracting the median value from each column. A two-sample unpaired student T-test was performed comparing the various sets of condition groupings (S_0_ = 0.1/0.5, FDR = 0.01/0.05) for all proteins remaining in the dataset and the results analyzed according to their statistical significance.

#### Proximity Based Labelling

6 x 10^6^ cells stably expressing TurboGFP clones 1 and 2 and ZDHHC20 clones with C-or N-terminally fused TurboID were plated in duplicate in 10 cm dishes. After reaching 80% confluency, cells were treated with 500 μM biotin for 3 h, cooled on ice, harvested and lysed in SDS lysis buffer (described above). A BCA assay was performed and two 1 mg portions of each lysate (n = 4 for each cell line) at 1 mg/mL were added to dimethylated neutravidin-agarose beads and agitated at RT for 3 h. Beads were washed 3X 0.2% SDS 50 mM HEPES pH 7.4 and 3X 50 mM HEPES pH 7.4. Beads were then suspended in 50 mM HEPES pH 8.0 containing 400 ng LysC for 1 h at 37°C. The supernatant was reduced with 5 mM TCEP and alkylated using 15 mM chloroacetamide for 15 min then digested with 100 ng of trypsin o/n at 37°C (Promega). Samples were acidified with 0.5% (v/v) trifluoroacetic acid (TFA) and the solvent removed in an Eppendorf Concentrator plus at 45°C.

After proteomes were labelled with TMT-10Plex reagents and combined solvent was removed in an Eppendorf Concentrator plus at 45°C. Samples were redissolved in 1% TFA and loaded on pre-activated 3 layers of SCX membranes and fractionated 6X. Membranes were washed 3X with 60 µL 0.2% TFA. Peptides were eluted using 6 elution buffers (60 µL); buffer 1 (75 mM ammonium formate, 20% (v/v) MeCN, 0.5% (v/v) formic acid), buffer 2 (125 mM ammonium formate, 20% (v/v) MeCN, 0.5% (v/v) formic acid), buffer 3 (200 mM ammonium formate, 20% (v/v) MeCN, 0.5% (v/v) formic acid), buffer 4 (300 mM ammonium formate, 20% (v/v) MeCN, 0.5% (v/v) formic acid), buffer 5 (400 mM ammonium formate, 20% (v/v) MeCN, 0.5% (v/v) formic acid), buffer 6 (5% (v/v) ammonium hydroxide, 80% (v/v) MeCN). Samples were dried in an Eppendorf Concentrator plus at 45°C and then dissolved in water containing 0.1% TFA, before analysis on a Q-Exactive mass spectrometer (Thermo Fisher) coupled to an Ultimate3000 LC (Thermo Fisher) using an Easy Spray Nano-source. The instrument was operated in data dependent acquisition mode selecting the 10 most intense precursor ions for fragmentation.

Data analysis was performed using Perseus (version 1.6.2.1). MaxQuant proteingroups.txt output files were uploaded and filtered against contaminants, reverse and proteins identified by site. A base 2 logarithm was applied to all reporter intensity corrected values and data filtered for where valid values were found in at least 8/10 channels. Data was normalized across all samples by subtracting the median across replicates within each TMT multiplex followed by normalizing across the conditions by subtracting the mean value from each column. A two-sample unpaired student T-test was performed comparing the various sets of condition groupings (S_0_ = 0.1, FDR = 0.01) for all proteins remaining in the dataset and the results analyzed according to their statistical significance.

#### Generation of ZDHHC20 knock-out cell lines

Two guide sequences (gRNA1 and 2) targeting exon 9 or 4, respectively, (Supplementary Table 7) were designed using the online tool CHOPCHOP (https://chopchop.cbu.uib.no)^57^ and separately cloned into a plasmid containing Cas9 and the sgRNA scaffold, pSpCas9(BB)-2A-Puro (PX459), using a Fast Digest BbsI restriction strategy coupled with T7 DNA ligase ligation. Plasmids were sequenced by GATC biotech to confirm subcloning of the gRNA guides. 1 μg of each plasmid was mixed with 250 μL of Opti-MEM before being combined with another mixture containing 5 μL of TransIT-X2 (5 µL/well) in 250 µL Opti-MEM. The combined mixtures were allowed to incubate at room temperature for 20 min before being added to a cell suspension containing 6 x 10^5^ HEK293T cells in a 6-well plate. The cells were culture for 3 days before selection in 1 μg/mL puromycin for 1 week. Cells were then single cell sorted into 96-well plates and allowed to expand into 12-well plates before screening by anti-ZDHHC20 and -calnexin (loading control) immunoblot. For single cell sorting, see **Molecular Cloning**.

#### Bioinformatics – PANTHER Overrepresentation Analysis

The online bioinformatic tool PANTHER^58^ was used to perform statistical overrepresentation analysis of ZDHHC20 substrates enriched in at least 2 of 3 cell lines: HEK293T, PANC1, and MDA-MB-231. Two separate analyses were performed using gene ontology terms cellular compartment and protein class. The cellular compartment (GO-Slim) analysis was performed using the default list of genes from the human genome, whereas the protein class analysis was done with a manually curated list representing the human S-acylated proteome. Statistical analysis and p-values were determined using an FDR adjusted Fisher’s Exact test. Results were filtered for those with a -Log_10_ (p-value) >9. The human S-acylome contains the combined unique hits between the following filtered SwissPalm lists: the first list was collated by setting the ‘nber_palmitoyl_proteome_hits’ and ‘nber_technique_categories’ ≥ 2 and the second list was generated by setting the ‘nber_palmitoyl_proteome_hits’ ≤ 1 and ‘nber_targeted_study_hits’ ≥ 1. After conversion of Uniprot AC IDs to gene name, the combined list of unique genes totalled 2429. Results were filtered for those with a -Log_10_ (p-value) >1.5.

## Data availability

The mass spectrometry proteomics data have been deposited to the ProteomeXchange Consortium via the PRIDE partner repository with the dataset identifier PXD032373 and PXD032378. Lipidomics datasets are currently under review at MetaboLights.

## Acknowledgements

**Acknowledgments**: The authors would like to thank Dr Brent Martin for engaging discussions on ZDHHC chemical genetics and Professors Yusuke Ohno and Akio Kihara from Hokkaido University for sharing their complete set of human N-terminal FLAG-tagged ZDHHC constructs. We would also like to thank Emmanuelle Thinon (Institut Européen de Chimie et Biologie, France) and Hans Bakker (Hannover Medical School, Germany) for generously gifting the Tate group with IFITM3, and VAMP3 and XXYLT1 plasmids, respectively. We thank Bram Snijders, Steven Howell, Vesela Encheva, and Joanna Kirkpatrick for their assistance and sharing of knowledge regarding LC-MS/MS as well as the Cell Services, Metabolomics, and the Flow Cytometry Science Technology Platforms at the Francis Crick Institute for their support and sharing of knowledge. Schematic figures were generated using BioRender.com.

## Funding

• CRUK/EPSRC Multidisciplinary Award to EWT and JD (C29637/A27506 and NS/A000078/1)

• CRUK Programme Foundation Award to EWT (C29637/A20183)

• CRUK Programme Award to EWT (DRCNPG-Nov21\100001), with support from the Engineering & Physical Sciences Research Council

• CRUK Convergence Science Centre studentship to ALL, EWT and JD (C24523/A27435)

• Wellcome Trust Senior Investigator Award awarded to JD (103799/Z/14/Z) and Wellcome Trust Investigator award (110060/Z/15/Z) to UE.

• Core funding from The Francis Crick Institute from Cancer Research UK (FC001070), the UK Medical Research Council (FC001070), and the Wellcome Trust (FC001070) was received by JD

• European Research Council Advanced Grant RASImmune awarded to JD

## Author Contributions

• Conceptualization: EWT

• Data Curation: CAO, MPB, JS, EMS

• Formal Analysis: CAO, MPB, JS, ALL, EMS, SAPD, JV

• Funding Acquisition: EWT, JD, CAO, USE

• Investigation: CAO, MPB, JS, ALL, EMS, SAPD

• Methodology: CAO, MPB, JS, ALL, EMS, JIM

• Project Administration: EWT, JD, USE

• Resources: CS, GT, JIM

• Software: EMS

• Supervision: EWT, JD, CAO

• Visualization: CAO, MPB, JS, EWT

• Writing – Original Draft Preparation: CAO, MPB, JS, ALL, EMS, EWT

• Writing – Review & Editing: All authors

**Competing interests**: EWT is a founder and shareholder in Myricx Pharma Ltd, and receives consultancy or research funding from Kura Oncology, Pfizer Ltd, Samsara Therapeutics, Myricx Pharma Ltd, MSD, Exscientia and Daiichi Sankyo. JD has acted as a consultant for AstraZeneca, Jubilant, Theras, BridgeBio, and Vividion and receives research funding from Bristol Myers Squibb and Revolution Medicines. All other authors declare no competing interests.

## Supplementary Information is available for this paper

Correspondence and requests for materials should be addressed to Ed Tate. Reprints and permissions information is available at www.nature.com/reprints.

